# gene-cocite: a web application for extracting, visualising and assessing the cocitations of a list of genes

**DOI:** 10.1101/109173

**Authors:** Richard Newton, Lorenz Wernisch

## Abstract

**Background:** The outcome from the analysis of high through-put genomics experiments is commonly a list of genes. The most basic measure of association is whether the genes in the list have ever been cocited together.

**Results:** The web application `gene-cocite` accepts a list of genes and returns a list of the papers which cocite any two or more of the genes. The proportion of the genes which are cocited with at least one other gene is given, and the *p*-value for the probability of this proportion of cocitations occurring by chance from a random list of genes of the same length calculated. An interactive graph with links to papers is displayed, showing how the genes in the list are related to each other by publications.

**Conclusions:** gene-cocite (http://sysbio.mrc-bsu.cam.ac.uk/gene-cocite) is designed to be an easy to use first step for biological researchers investigating the background of their list of genes.

## Background

The outcome from the analysis of high through-put genomics experiments is commonly a list of genes. Often the next step in the analysis is to look for known connections between the genes using some form of geneset enrichment analysis, where the content of the gene list is compared to lists of genes already known to be associated in some way [1, 2], for example within a biological pathway or GO annotation [3].

A more basic measure of association is whether the genes in the list have ever been cocited together within publications. The web application `gene-cocite` accepts a list of genes and returns a list of the papers which cocite any two or more of the genes. The proportion of the genes in the list which are cocited with at least one other gene is also given. And the *p* -value for the probability of this proportion of cocitations occurring by chance from a random list of genes of the same length is calculated. An interactive graph with links to papers is displayed, showing how the genes in the list are related to each other by publications. `gene-cocite` (http://sysbio.mrc-bsu.cam.ac.uk/gene-cocite) is designed to be an easy to use first step for biological researchers investigating the background of their list of genes.

## Implementation

The source of the information on genes and publications used by `gene-cocite` is PubMed. PubMed [4] contains information on more than 24 million papers, including details on which genes each of the articles cite. Each article in PubMed has a unique identifier, the PubMed ID. The underlying analysis of `gene-cocite` is performed using R [5]; specifically packages from the Bioconductor project [6]. The R package org.Hs.eg.db [7] (and fifteen similar packages for other organisms) contains mappings between gene identifiers and PubMed IDs. For each gene identifier the package provides a list of the PubMed IDs of the papers which cite this gene. The packages are updated every 6 months and the versions being used currently by `gene-cocite` are listed in a link from the main web page. Using the gene identifier to PubMed ID mappings, cocitations can be easily extracted.

The proportion of the gene list which cocite with at least one other gene in the list is calculated. The *p* -value of obtaining a proportion p of cocited genes from a gene list of length *n* is obtained by taking *m* random samples of *n* genes from a list of all the genes in the database and recording for each sample what proportion *p^*^* are cocited together in at least one paper. The *p* -value is then the fraction of *p\** greater or equal to proportion *p*. For gene lists of length 2 to 400 the random samples are pre-calculated with *m* = 10000. For gene lists longer than 400, random samples have been calculated at increments of 100 genes up to lists of length 6000 genes. Since the distribution of the random samples for gene lists of this length is close to being normal, the mean and standard deviation of the sample at each increment are calculated. Then to predict results for list sizes intermediate between the increments, two loess curves are fitted, one through the mean values for all increments of gene list length and one through the corresponding standard deviations. The two loess curves are then used to predict the mean and standard deviation of the normal distribution of the random sample for any gene list of length greater than 400. The normal distribution is then used to calculate the *p* -value for a given proportion of genes cocited.

The basic framework of the web application is constructed using Apache Struts, which itself uses JavaServer Pages (JSP) and Java, running on an Apache Tomcat server connected to the outside world via an Apache HTTP server. Information on how a Struts web application can be made to run an R script and return the results to the user’s web page can be found in [8].

The output from the R script, namely the summary information, the table of gene pairs with their citing papers and the data for the interactive graph, is returned to the web page as JavaScript Object Notation (JSON) files. These are processed on the web page using JavaScript. The sortable table is created using the JavaScript package Tidy Table [9] and the interactive force layout graph showing how genes are connected by publications is produced using the JavaScript library D3 [10].

## Results and Discussion

Gene lists can be entered as either HUGO [11] symbols or as Entrez Gene IDs [12] The databases that `gene-cocite` utilises contain information on sixteen different organisms: Anopheles (*Anopheles gambiae*), Bovine (*Bos taurus*), Canine (*Canis familiaris*), Chicken (*Gallus gallus*), Chimp (*Pan troglodytes*), *Escherichia coli*) (*strain K12*), *Escherichia coli*) (strain Sakai), Fly (*Drosophila melanogaster*), Human (*Homo sapiens*), Mouse (*Mus musculus*), Pig (*Sus scrofa*), Rat (*Rattus norvegicus*), Rhesus (*Macaca mulatta*), Worm (*Caenorhabditis elegans*), Xenopus (*Xenopus laevis*) and Zebrafish (*Danio rerio*). The organism of interest is selected on the web page. The only other input required from the user is the gene citation threshold. In performing a cocitation analysis it is usually advisable to restrict the search to papers that cite just a small number of genes. Papers which cite many thousands of genes will not be particularly informative, so are best excluded from the analysis. `gene-cocite` allows the user to restrict the search to papers which cite a maximum of 10, 50, 100, 250 and 500 genes.

The web page displays a synopsis of the analysis, namely the number of genes analysed, the number of publications found, the percentage of genes which are cocited with at least one other gene in the list and the *p* -value for this proportion of cocitations (Figure 1). A significant *p* -value suggests the corpus of scientific publications has revealed an association of the genes. Given that little is known about the vast majority of genes and their interactions, a non-significant result may simply indicate that some or all of the genes in list are still poorly categorized.

**Figure 1.**
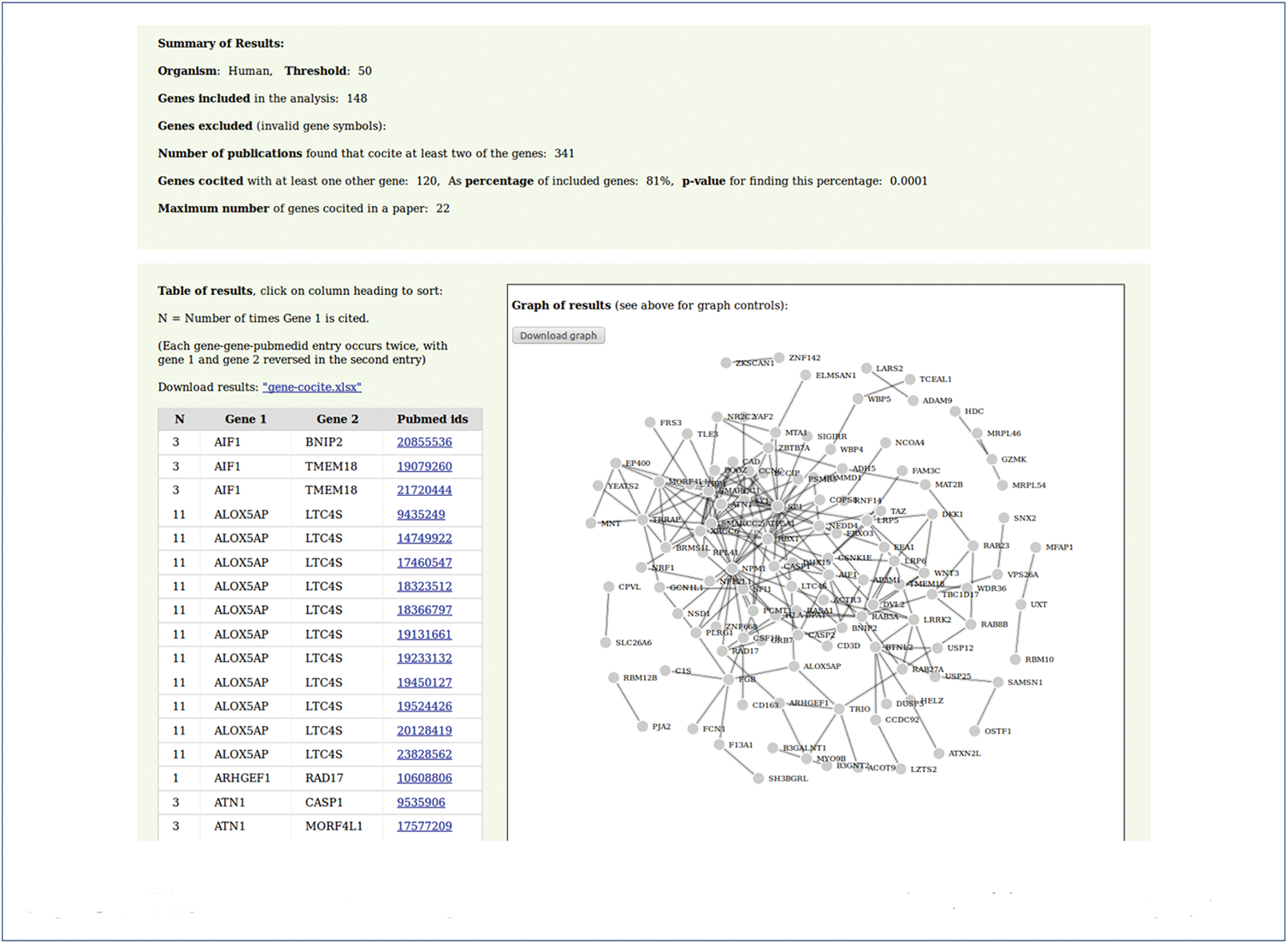
The results section of gene-cocite. The summary results, table and interactive graph

Below the summary of results, the detailed results are presented in two different formats. On the left of the screen, the results are shown as a sortable table giving gene pairs and the publication which cocites them (as a link to a PubMed entry). And on the right of the screen the results are presented as an interactive graph that uses the D3 [10] force layout. Once the force layout has reached a stable network configuration, hovering on a node of the graph will highlight the corresponding gene’s nearest neighbours, that is, all the genes which are cocited with the gene (Figure 2). Clicking on the node isolates this sub-network (Figure 3). Clicking on a network edge will display a tooltip with the PubMed ID (or IDs) of the article(s) which cocite the two genes connected by the edge (Figure 4). As in the table, the PubMed IDs are presented as links to the PubMed entries for the papers. Both the results as a table (as a .xls file) and the results as a graph (as a .png file) are downloadable by the user.

**Figure 2.**
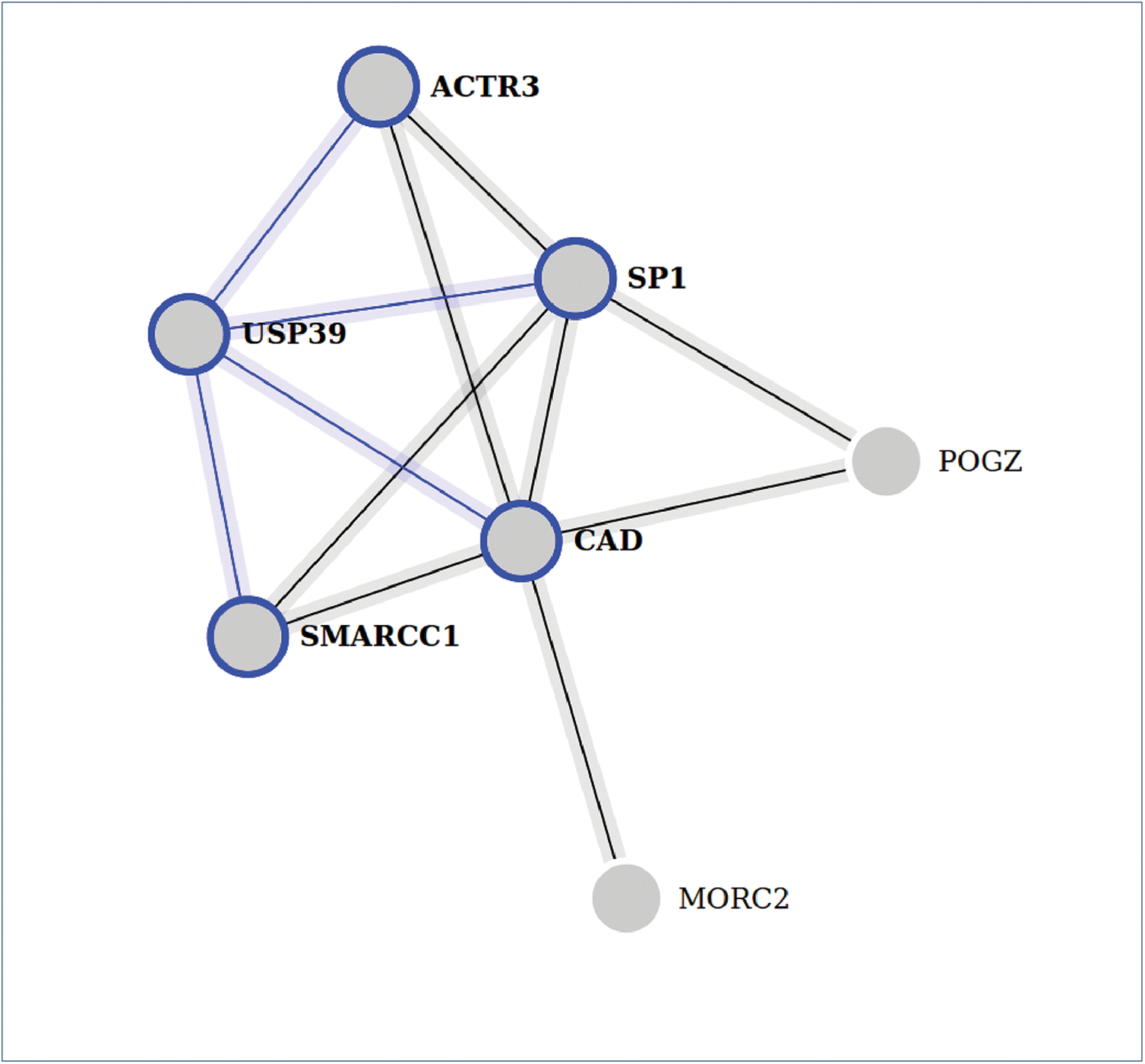
Hovering over a gene node. Hovering over a gene node highlights the gene's nearest neighbours

**Figure 3.**
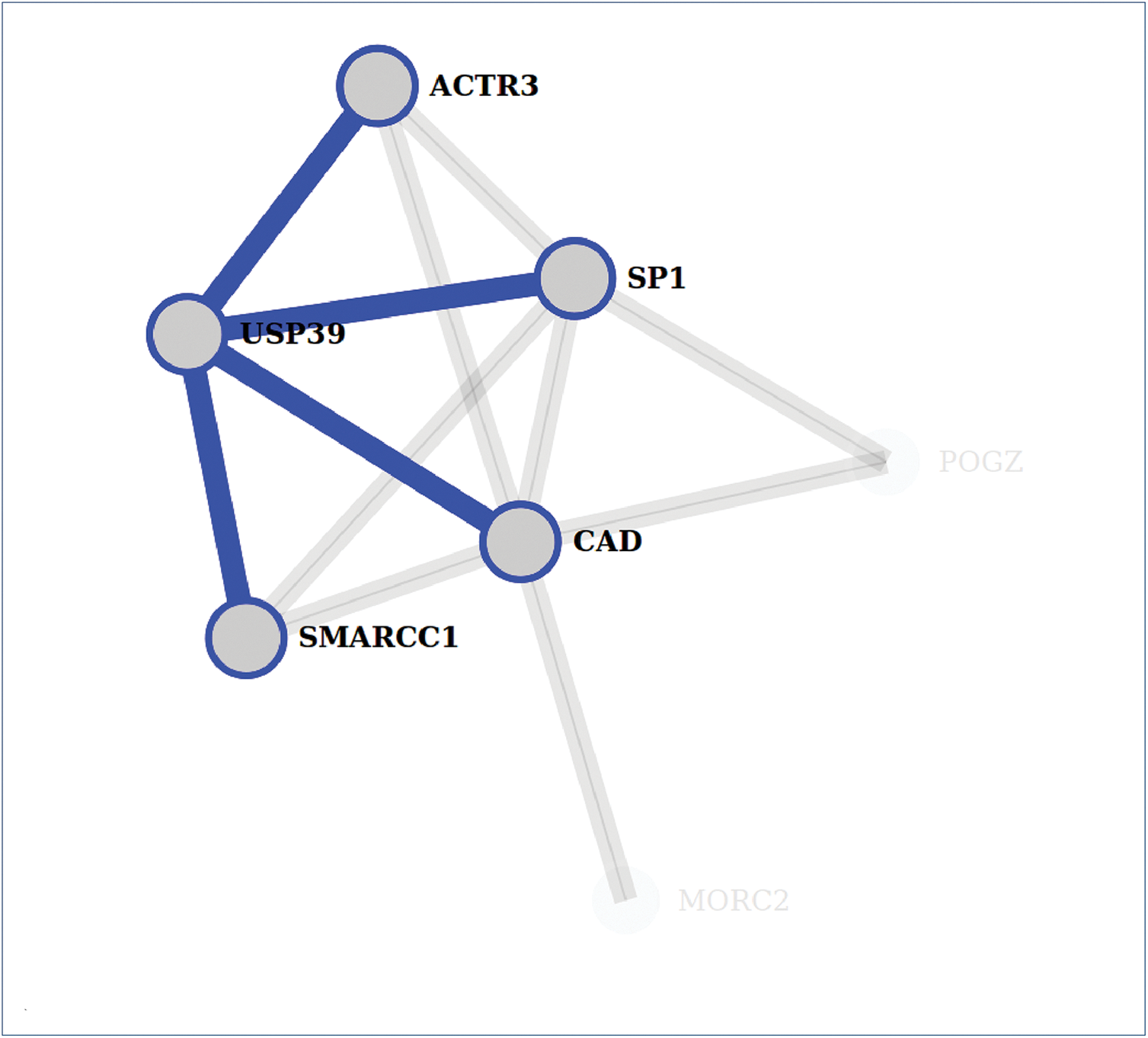
Clicking on a gene node. Clicking on a gene node isolates the gene's sub-network

**Figure 4.**
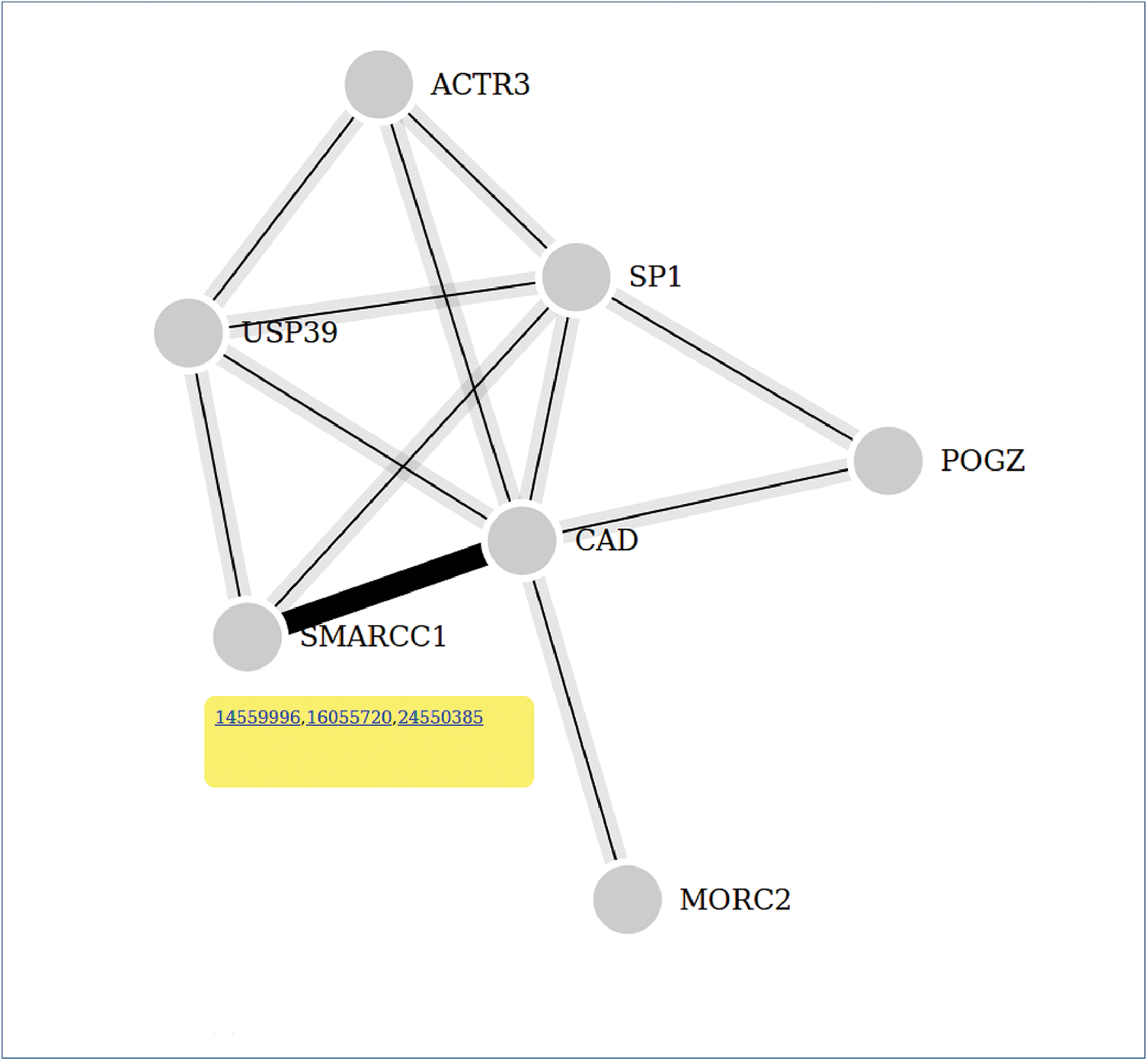
Clicking on an edge. Clicking on an edge displays a tooltip with the PubMed IDs of the articles which cocite the two genes connected by the edge

The information generated by `gene-cocite` could be found through a direct use of R, but `gene-cocite` is designed for ease of use by those unfamiliar with R. A gene list can be pasted directly into the search page at PubMed and this would produce a list of citing papers. However papers that cite just one of the genes in the list would be included in the output and it would also require some effort to identify exactly which combinations of genes are being cocited in which papers.

The extensive web application Coremine medical [13] can generate cocitation results from gene lists (comprising 300 or fewer genes). Coremine medical is a versatile tool incorporating far more sources of information, but with a concomitant increase in the complexity of use and the visual complexity of the results presented to the user. `gene-cocite` is simpler in its aims and scope and hence simpler in its usage. A comprehensive comparison has not been made but `gene-cocite` does currently appear to identify more publications than Coremine medical. Text mining based applications such as Cociter [14] take gene lists as input and utilise cocitations, but in the context of making functional assignments for the genes. `gene-cocite` is not intended as a tool for automatically extracting biological function, but to simplify a cocitation search, present results to the user in an easily accessible form and to provide an indication of the probability of finding this number of cocitations by chance.

## Conclusions

`gene-cocite` is intended as a simple to use tool for the preliminary analysis of gene lists; as a step between browsing PubMed and looking for functional assignments. Sixteen different organisms can be investigated. It presents the papers which cocite any of the genes contained in the user supplied list in both a tabular and a graphical form. And it assess whether, based on the cocitations, the association between the genes in the list is more than would be expected from a randomly chosen list of genes of the same length. The databases used by `gene-cocite` will be updated as and when new versions of the Bioconductor packages from which they are derived become available.

## 1 Availability and requirements

- Project name: `gene-cocite`
- Project home page: http://sysbio.mrc-bsu.cam.ac.uk/gene-cocite
- Operating system(s): Platform independent
- Programming language: R, Java, JavaScript, D3
- Other requirements: Up to data browser - tested on Mozilla Firefox version 40.03, Chrome version 45.0 and Internet Explorer version 11.0.22. (The D3 generated graph will not be visible in Internet Explorer 8 or less). Depending on the number of nodes in the graph the D3 force layout can be quite demanding on processing power whilst searching for an equilibrium, so on older machines this may take some time.
- License: No license
- Any restrictions to use by non-academics: None

## Competing interests

The authors declare that they have no competing interests.

## Author’s contributions

The authors contributed equally to the publication.

